# Protocol for green synthesis of excellent fluorescent mango leaves-derived carbon for advanced theragnostic in biological applications

**DOI:** 10.1101/2025.10.28.685050

**Authors:** Ankesh Kumar, Himanshu Khatri, Raghu Solanki, Pankaj Yadav, Dhiraj Bhatia, Amit K. Yadav

## Abstract

Carbon quantum dots (CQDs) are zero-dimensional fluorescence nanoparticles that are less than 10 nm in size. They have unique properties like tunable surface functionality, biocompatibility, fluorescence, and low toxicity. This study outlines a detailed protocol for synthesizing CQDs from mango leaf powder using a reflux method at 160 C and 400 rpm for a duration of two hours. The synthesized CQDs were comprehensively characterized using the following techniques, including UV-visible spectroscopy, fluorescent spectrophotometry, atomic force microscopy, dynamic light scattering and zeta potential measurement, Fourier transform infrared spectroscopy, and scanning electron microscopy. In vitro analysis was conducted through an MTT assay, revealing that the CQDs exhibit reduced toxicity in RPE1 cells compared to the control group. Additionally, a concentration-dependent cellular uptake assay was performed to further evaluate the biological performance of the CQDs. The findings suggest the potential of CQDs as safe and effective nanomaterials for various biological applications, including drug delivery and bioimaging.

## 1. Introduction

Carbon quantum dots are zero-dimensional nanoparticles with dimensions typically below 10 nm. These carbon dots have unique properties that include size, fluorescence^1^, and surface morphology, tunable surface functionality^2^, photostability, low toxicity^3,4^, water solubility and biocompatibility^5^. These attributes make them useful to various applications, including drug delivery^6^, bioimaging^7^, phototherapy^8^, energy storage systems^9^, and antimicrobial protection agents^10^. Additionally, these are used as non-material-based formulation applications in wound healing due to their antimicrobial^11,12^ and anti-inflammatory properties^13^.

Several scientists have synthesized carbon quantum dots to improve fluorescence by utilizing chemical-based precursors. These carbon dots show toxicity. In this paper, carbon quantum dots (CQDs) were synthesized using natural sources^1,3,13–15^.

Photoluminescent CQDs were synthesized by utilizing bottom-up approaches involving ultrasonication, microwave irradiation, hydrothermal/solvothermal, and the reflux method^16^. Green-synthesized CQDs have good biocompatibility and excellent fluorescence^17^. In addition, the CQDs were synthesized using different types of natural precursor resources, such as green tea leaves, orange peel, garlic, and other bio-waste for CQDs synthesis^18^. These natural precursors help to overcome surface passivation. Therefore, careful selection of precursor materials is essential to understand their biological, physical importance and green synthesis^7^. The medicinal plant should be the most suitable precursor for CQDs synthesis. Biowaste materials have gained prominence in the production of fluorescent CQDs due to their eco-friendliness, easy synthesis, low cost, therapeutic properties, biocompatibility, and bioavailability^19^.

In this study, we synthesized CQDs from fresh green mango leaves (*Mangifera indica*). The synthesis process involved using an ethanolic extract of the leaves, which was prepared using the reflux method^20,21^. This technique operates as a closed system, maintaining a constant temperature throughout the reaction, continuous stirring, and preventing any loss of solvent. In addition, the mango powder is mixed with ethanol. The resulting ethanolic extract initiates the reaction, starting the carbonization process. Carbonization is the process by which the precursor material is reduced to the nanoscale using temperature and mechanical methods. This method is easy to set up and forms uniform nano-sized CQDs with high yield^22^.

The highly red fluorescent CQDs are synthesized from mango leaves. The mango plant (Scientific name: *Mangifera indica)*^2,3^. Mango leaves are valued for their medicinal properties and are frequently utilized in traditional therapies, particularly in Ayurveda and traditional Chinese medicine^23^. They are abundant in beneficial compounds such as phenolic acid, polyphenols, flavonoids (including quercetin), vitamins A, B, C, and E, as well as essential minerals like potassium, calcium, and magnesium. These components bestow mango leaves with antioxidant and anti-inflammatory effects, helping protect cells and reduce inflammation^24^. Research indicates that they may assist in regulating blood sugar levels, preventing diabetes-related complications, and promoting cardiovascular health, while also exhibiting antimicrobial activity and potential anti-cancer properties. Traditionally, mango leaves have been used to address digestive issues, respiratory conditions such as bronchitis and asthma, and to support wound healing. However, more clinical studies are necessary to establish safe and effective dosages for specific conditions before they can be routinely recommended for therapeutic use^1,25–32^.

There are commercially available fluorophores that face challenges due to their low cellular permeability. To address these issues, the use of smaller, more prominent organic fluorophores with good surface functional groups helps enhance their ability to permeate within cells. Cell-impermeable substances like drugs, proteins, and nucleic acids can enter live cells through simple incubation^33^. Consequently, cell-penetrating peptides provide an effective method for delivering nonpermeable organic fluorescence probes into cells^1,2,7,21,27^. The CQDs are used as fluorescent probes for bioimaging and are also used in drug delivery, owing to their biocompatibility, small size, and low cytotoxicity^32^. In 2025, studies demonstrated the versatility of carbon dots (CDs) in biomedical applications. A key advancement was the creation of a red-emitting carbon dot composite (CDs@BSA) for targeted bioimaging in rheumatoid arthritis, facilitating real-time monitoring^34^. Researchers also developed biocompatible blue-fluorescent CDs from PEC-GS/BG hybrids, effective for stable cell imaging^35^. Moreover, multifunctional CD protein conjugates, like CD–pepsin nanoparticles created using machine learning, improved immune cell imaging and offered sustained drug delivery and photodynamic therapy^36^. Another study produced carbon dots from citric acid and o-phenylenediamine, achieving strong fluorescence suitable for organelle-specific imaging in plants and animals^37^. Carbon dots provide excellent optical stability, photobleaching resistance, and customizable surface functionalization, making them applicable for imaging, diagnostics, and therapy integration^18^.

This study presents a novel approach to the green synthesis of carbon quantum dots (CQDs) utilizing mango leaf powder through a reflux method. The synthesized CQDs were characterized through a comprehensive array of techniques. Optical properties were evaluated via UV-Vis spectroscopy and fluorescence spectrophotometry. Size distribution and surface charges were determined using Dynamic Light Scattering (DLS) and Zeta potential measurements, respectively. The functional groups on the surface of the CQDs were analyzed using Fourier Transform Infrared (FTIR) spectroscopy. Structural characterization was accomplished through X-ray Diffraction (XRD), while the morphology and shape of the CQDs were investigated using Atomic Force Microscopy (AFM) and Transmission Electron Microscopy (TEM). Additionally, cellular interactions were assessed through MTT assays and cellular uptake studies to evaluate the CQDs’ biocompatibility and efficacy for potential applications in biological systems.

## 2. Overview of the procedure

### 2.1 Precursor material

Mango leaves, used in traditional therapies like Ayurveda and Chinese medicine, are rich in beneficial compounds such as phenolic acids, polyphenols, flavonoids (including quercetin), and vitamins A, B, C, and E. They also contain essential minerals like potassium, calcium, and magnesium, which provide antioxidant and anti-inflammatory effects, protecting cells and reducing inflammation^24^.

### 2.2 Reflux synthesis method to produce carbon quantum dots (CQDs) derived from mango leaves (mQDs)

The reflux method heats a reaction mixture to its boiling point while condensing the vapors back into liquid. This is done by attaching a condenser to the reaction vessel, which cools the vapor using chilled water, maintaining a closed system and constant solvent volume. This technique enables efficient reactions that require prolonged heating without solvent loss to evaporation, maintaining a steady reaction temperature at the solvent’s boiling point^38^.

### 2.3 Comparison with other methods

Carbon Quantum Dots can be synthesized using various methods, with the reflux method standing out for its simplicity and practicality. In this approach, reaction precursors are dissolved in a solvent and heated under reflux, using a condenser to prevent solvent loss. This technique enables effective control over particle growth, resulting in CQDs that are generally uniform, biocompatible, and water-soluble. The main advantages of the reflux method include minimal equipment requirements, cost-effectiveness, suitability for large-scale synthesis, and the ability to produce QDs with properties conducive to biological and imaging applications. However, this method typically requires longer reaction times compared to some rapid synthesis techniques. Additionally, it may offer less precise size control and lower quantum yields than more advanced methods like hot injections. In comparison, alternative techniques such as hot injection can yield very high-quality, monodisperse quantum dots with narrow size distributions and high quantum yields. However, these methods demand precise temperature control, specialized equipment, and are less practical for scaling up^38–40^. Other synthesis techniques, like sol-gel, hydrothermal/solvothermal, and ultrasonic chemical synthesis, each present unique advantages regarding property tuning and speed, but they also have limitations concerning equipment complexity, scalability, or quantum efficiency.

Overall, the reflux method finds a valuable balance among practical feasibility, cost, and product quality, making it an excellent choice for many research and biomedical applications, even though other methods may be preferred when the absolute highest optical performance is required^37^.

### 2.4 Application of the methods

Carbon Quantum dots are generally used in several biological applications that involve Drug delivery, bioimaging, and biosensing. Due to CQDs having excellent properties, for example, small size, good surface functionality, biocompatibility, and excellent photoluminescence^10,18,27^.

#### Bioimaging

Commercial fluorescent dyes like propidium iodide and propidium azide are effective at penetrating the cell walls of dead bacteria, which makes them popular choices for DNA staining methods. However, their high cost, toxicity, and tendency to undergo photobleaching are prompting researchers to look for dyes that are more affordable, less toxic, and highly soluble in water for cell viability assays. In contrast to conventional organic dyes, CQDs exhibit a broad spectrum of fluorescent properties that could be applied across various biological fields ^2,7,19,21,36^.

#### Drug delivery

In contrast to earlier metallic or inorganic nanoparticles, carbon dot delivery systems may lower cytotoxicity and enhance clinical results ^21^. Furthermore, their size is under 10 nm, which makes them ideal for drug delivery purposes ^6,36^.

#### Biosensing

Carbon Quantum Dots (CQDs), an advanced category of fluorescent nanomaterials, exhibit considerable promise for biosensing applications, owing to their superior photostability, biocompatibility, and tunable fluorescence properties. These nanostructures can be employed for the detection of biomolecules such as proteins, DNA, and various pathogens. By functionalizing CQDs with specific ligands, researchers can significantly improve both the sensitivity and selectivity of diagnostic assays. Their capability for rapid, real-time analysis positions them as valuable tools in healthcare diagnostics, environmental monitoring, and food safety assessments, facilitating the development of innovative and efficient sensing technologies ^41^.

### 2.5 Limitations of the protocol

The synthesized carbon dots face several limitations that impact their practicality. Firstly, the production yield is relatively low, generating only 25 mg per synthesis, which may not be adequate for larger applications. Additionally, these carbon dots demonstrate poor solubility in water and other common solvents, restricting their usability in various formulations. While they dissolve easily in ethanol, this reliance on a specific solvent could limit their compatibility in scenarios requiring aqueous solutions. Furthermore, the carbon dots must be stored in a dark environment to preserve their stability, presenting potential challenges for long-term storage and usage in diverse settings^20,21,38–40^.

## 3. Experiment design

### 3.1 Synthesis of mango leaves-derived quantum dots (mQDs)

Mango leaves provide an eco-friendly source for synthesizing mQDs, which are beneficial for bioimaging and cellular uptake. They can be synthesized using simple methods like reflux Techniques, resulting in highly fluorescent and biocompatible QDs. Research shows that QDs have excellent cellular uptake with low toxicity while exhibiting red or near-infrared fluorescence, ideal for bioimaging. They can penetrate cell membranes without affecting cell viability, making them suitable for intracellular imaging. When coated with specific cationic lipids, these quantum dots enhance fluorescence intensity and stability. Additionally, mango leaves contain natural antioxidants that improve the biological safety of CDs compared to traditional heavy-metal quantum dots. A study in Nature highlighted their potential in advanced bioimaging and photonic applications, showcasing their value in biomedical research and cellular imaging^20,21^.

### 3.2 Cytotoxicity

Cytotoxicity studies utilizing the MTT assay focus on evaluating the effects of carbon dots on cell viability. The procedure begins with seeding cells into 96-well plates, allowing them to adhere and settle properly. The cells are then treated with graphene or green carbon dots for 24 hours to assess their potential impact on viability. After treatment, MTT reagent is added to each well, followed by a 4-hour incubation period. During this time, viable cells convert the yellow MTT compound into insoluble purple formazan crystals, facilitated by the action of mitochondrial reductase enzymes. Following this incubation, dimethyl sulfoxide (DMSO) is introduced and allowed to incubate for 10 minutes to dissolve the formazan crystals, making the solution suitable for spectrophotometric detection. Finally, the absorbance of the resulting solution is measured at 570 nm, providing a quantitative assessment of cell viability or cytotoxicity based on the metabolic activity of the cells. This procedure allows researchers to efficiently evaluate the cytotoxic effects of carbon dot nanoparticles on cultured cells^42^.

### 3.3 Mechanism of cellular uptake of mQDs

Carbon dots are introduced into a cell, and they can be internalised through several endocytic pathways, including micropinocytosis, clathrin-mediated endocytosis^43^, and caveolae-mediated endocytosis^44^. In micropinocytosis, the cell membrane engulfs extracellular fluid and carbon dots, forming a macropinosome. This vesicle often fuses with a lysosome, leading to the degradation of the carbon dots within the lysosome. In clathrin-mediated endocytosis, carbon dots are internalized within clathrin-coated vesicles, which mature into late endosomes before fusing with lysosomes^28,43^. This process also results in the degradation of nanoparticles. Conversely, caveolae-mediated endocytosis employs specialized, flask-shaped membrane invaginations to create caveolae-coated vesicles containing carbon dots^22^. These vesicles develop into caveosomes, which can directly deliver their cargo to the endoplasmic reticulum (ER), effectively bypassing lysosomal degradation. The specific pathway taken by carbon dots significantly influences their intracellular fate. Caveolae-mediated uptake provides a route to functional cellular compartments like the ER, while macropinocytosis and clathrin-mediated endocytosis primarily direct the carbon dots towards lysosomal degradation. This distinction has important implications for designing carbon dot-based bioimaging agents or drug delivery systems, as the chosen endocytic route affects the stability and efficacy of these particles within biological cells^5,22,43^.

## 4. Materials

### 4.1 General Solvent

- Ultrapure water (Milli-Q water)
- Dimethyl sulfoxide
- Ethanol, 70% (vol/vol)
- Phosphate-buffered saline (PBS), pH 7.4
- Mango leaves fine powder from the IIT GN campus mango tree

### 4.2 Cell culture

- Fetal bovine serum (FBS; Gibco)
- Dulbecco’s modified Eagle medium (DMEM) (Thermo Fisher)
- Penstrap(Gibco)
- 4′,6-Diamidino-2-phenylindole (DAPI; Roche)

Caution – DAPI is toxic. Wear appropriate personal protective equipment.

- Mowiol
- Trypsin–EDTA, phenol red (Thermo Fisher Scientific)
- HAMS-F12 media (Thermo Fisher)
- MTT Salt
- 0.1% Triton X-100
- 4% paraformaldehyde
- 0.4X Phalloidin A488 (Sigma Adrich)

### 4.3 Equipment

- Multichannel pipettes (Eppendorf)
- Pipette(Eppendorf)
- Pipette tips(Tarsons)
- Vortex mixer (Thermo Scientific)
- Water bath (Scientz)
- Ice machine (Coolium)
- Refrigerated benchtop centrifuge (Eppendorf)
- Freezers and refrigerators (Thermo Fischer)
- pH meter (Mettler Toledo)
- Bath Sonicator (Cole-Parmer)
- Ultra-pure water purifier (Milli-Q)
- Autoclave (Bionexis)
- Filter, 0.22-μm polyethersulfone membrane (Avantor)
- Analytical balance (Lab science)
- Confocal microscope (Leica Microsystems, Germany)
- Fluorescence spectrophotometer (FP-8300 Jasco spectrophotometers (Japan))
- Dynamic light scattering (Malvern Panalytical Zetasizer Nano ZS)
- UV-VIS/fluorescence Cuvettes (Standard Cell with Lid & round Bottom, Dimensions:45×12.5×12.5)
- Centrifuge (Remi Bench Top Refrigerated Centrifuge)
- UV-visible spectrophotometer (Spectrocord-210 Plus Analytokjena (Germany))
- Atomic force microscopy (Bruker)
- Mica substrate
- Wash bottle(100ml) (borosil)
- Centrifuge tubes (Falcon 15 mL and 50 mL centrifuge tubes, Tarsons, Cat no.546041)
- FTIR Spectrometer (PerkinElmer)
- Parafilm tape(Amcor)
- Glass vial(10ml) (Borosilicate)
- Syringe needle (Dispo van)
- Rotary evaporator (Heidolph 036040051 Rotary Evaporator, Manual, G3, Std Distillation Vertical Condenser)
- Desiccator (PolyLab Plastic Vacuum)
- Reflux condenser (Borosilicate)
- Microscope slide (Bluestar)

### 4.4 Cell culture

- Hemo cytometer (Thermo Fisher Scientific)
- Cell culture flask, 25 cm^2^ (NEST)
- Cell culture dish, 10 cm (NEST)
- 10 mm glass-bottomed Petri dish (NEST)
- Cell culture incubator (Thermo Fisher Scientific)
- Cell culture microscope (Carl Zeiss)
- Serological pipettes (Tarsons)
- 96-well plate (Sigma Aldrich)
- 4-well plate cell culture(Sigma Aldrich)

### 4.5 Software

- ImageJ software
- Graph paid
- Origen 2025

## 5. Experimental setup

The experimental setup requires the following equipment: a hot plate magnetic stirrer, a silicon oil bath, a 100 ml two-round-bottom flask, a water inlet and outlet pipe, a condenser, a water pump, a fume hood, and a bucket of ice water. The reflux condenser contains mango leaf extract and is connected to a magnetic bead-filled round-bottom flask (RBF) using a Teflon tap. The silicon oil bath is placed on the hot plate magnetic stirrer, and the condenser and RBF assembly are secured in the silicon oil bath using a clamp. The inlet and outlet of the condenser are connected to the water pump with the appropriate piping.

The glassware employed within this protocol must be cleaned with aqua regia and deionized water as many times as possible to eliminate substances on its surface. Aqua regia is a toxic mixture of acids that may cause severe injuries to the skin or organs in the event of inhalation or contact. Wear the correct protective laboratory equipment, such as gloves, mask, laboratory coat, and goggles, and handle the aqua regia in a fume hood.

It is important to notice that the round-bottomed flask should not contain more than half of the volume of the reaction liquid.

## 6. Reagent Preparation

### 6.1. Mango Extract

2 grams of mango leaf powder were supplemented in 30 ml of solution (composed of 20 ml of ethanol and 10 ml of Milli-Q water). The solution bottle was continuously mixed for 4 hours. Afterward, the extract was collected and transferred into a Falcon tube, where it was centrifuged. The pellet was discarded, and the supernatant containing the mango leaf extract was collected.

### 6.2 Aqua regia

Aqua regia is composed of concentrated solutions of nitric acid (HNO3, 65 wt%) and hydrochloric acid (HCl, 30 wt%) in a molar ratio of 1:3 (HNO3/HCl). It is essential to mix the HNO3 and HCl solutions carefully. Always handle aqua regia with caution, wearing gloves and goggles for safety. Since aqua regia quickly loses its effectiveness, it is important to prepare a fresh solution each time it is needed. Additionally, keep aqua regia in an open container, as it will degrade over time and produce harmful gases.

### 6.3 DMEM

To 500 ml DMEM, add 8% (vol/vol) FBS and 5 ml 100X penicillin-streptomycin solution. The medium may be frozen up to 2 months at 4 °C. Warm it to 37 °C before usage.

### 6.4 HAMS-F12 media

F-12 medium consists of several key components, including zinc, putrescine, hypoxanthine, and thymidine, but is devoid of proteins and growth factors. Consequently, it is typically supplemented with 10% fetal bovine serum (FBS) to enhance its nutritive capacity. The medium utilizes a sodium bicarbonate buffer at a concentration of 1.176 g/L, which necessitates a CO2 environment maintained at 5–10% to uphold physiological pH levels.

### 6.5 1X Phosphate-buffered saline (PBS)

To prepare a phosphate-buffered saline (PBS) solution, accurately weigh and dissolve the following reagents in 800 mL of deionized water: 8.0 g of sodium chloride (NaCl), 0.2 g of potassium chloride (KCl), 1.44 g of disodium hydrogen phosphate (Na_2_HPO_4_), and 0.24 g of potassium dihydrogen phosphate (KH2PO4). Adjust the pH to 7.4 using hydrochloric acid (HCl) as necessary. Once the pH is stable, bring the total volume up to 1,000 ml with deionized water. Subsequently, sterilize the solution by passing it through a 0.22 µm membrane filter. Store the resulting PBS solution at 4°C, where it can remain stable for up to six months.

### 6.6 MTT Solution

To prepare a 15 ml stock solution of MTT salt at a concentration of 5 mg/ml, you need to dissolve 75 mg of MTT salt into 15 ml of 1x PBS buffer.

For the working solution, which requires a concentration of 0.5 mg/ml with a total volume of 10 ml in serum-free media, take 1 ml of the stock solution and dissolve it in 9 ml of serum-free media. This will result in a prepared working solution of 0.5 mg/ml.\

### 6.7. 4% paraformaldehyde

To prepare a 4% Paraformaldehyde solution, start by measuring 5.4 mL of a 37% Paraformaldehyde stock solution. Next, dissolve this measured stock solution in 44.6 mL of 1X PBS (Phosphate-Buffered Saline). It is important to mix the solution thoroughly to ensure it is homogeneous. Once prepared, the resulting solution will have a final concentration of 4% Paraformaldehyde. For optimal results, store the solution at 4°C and use it within a week.

### 6.8. 0.1% Triton X-100

To prepare a 0.1% Triton X-100 solution, start by measuring 50 µL of Triton X-100. Next, add this volume to 49950µL of 1x PBS (phosphate-buffered saline). Ensure that you mix the solution thoroughly to achieve complete dissolution of the Triton X-100 in the PBS. Once mixed, the solution is ready for use in your experiments.

### 6.7. 0.4X Phalloidin A488

Prepare a 0.4X phalloidin solution by dissolving 50 µL of the 400X stock solution in 49950 µL of 1X PBS. This will result in a total volume of 50 mL of 0.4X phalloidin solution A488.

### 6.8. Mowiol DAPI

**-**In 1 mL of Mowiol, add 1 µL of DAPI.

## 7. Procedure

### 7.1 Mango leaves powder preparation

1.1. Gather mango leaves from a mango tree situated at the IIT Gandhinagar campus.

1.2. Washing Procedure: Rinse mango leaves with tap water, followed by a rinse using Milli-Q water.

1.3. Drying the Leaves: Wipe each leaf with tissue paper to remove any residual moisture, ensuring no tissue residues or dust are left on the surface. Dry the leaves in a closed, dust-free environment for approximately 10 to 12 days.

1.4. Grinding: Once dried, grind the leaves in a mixture grinder until a fine powder is obtained.

Ensure that the grinder is properly sterilized and clean.

1.5. Storage: Store the resulting mango leaf powder in 50 ml sterile Falcon tubes., Seal the tubes with parafilm and keep them at room temperature.

### 7.2. Glassware Cleaning for the Synthesis of CDs

- 2.1. Gather Required Glassware:

- Two 100 ml round-bottom flasks
- One condenser
- 2.2. Gather Additional Equipment:

- Magnetic stirrer
- Hot plate
- Silicon oil bath
- Water pump
- Ice water bucket
- 2.3. Prepare Aqua Regia Solution:

- Prepare and mix 180 ml of aqua regia by using a 3:1 ratio of hydrochloric acid to nitric acid.
- 2.4. Fill Glassware:

- Transfer the prepared aqua regia solution into the glassware according to its volume.
- 2.5. Fume Hood:

- Place the filled glassware in a fume hood chamber and let it sit for 24 hours.
- 2.6. Neutralize Aqua Regia:

- After 24 hours, neutralize the aqua regia solution using sodium hydroxide pellets.
- 2.7. Dispose of Neutralized Solution:

- Carefully dispose of the neutralized solution in the sink with flowing water.
- 2.8. Wear Safety Gear:

- Ensure to wear a proper lab coat, gloves, and goggles throughout the procedure.
- 2.9. Initial Cleaning:

- Wash the glassware with detergent and tap water.
- 2.10. Rinse with Milli-Q Water:

- Rinse the glassware thoroughly with Milli-Q water.
- 2.11. Wash with Acetone:

- Wash the glassware with acetone, followed by another rinse with Milli-Q water.
- 2.12. Dry in Hot Air Oven:

- Place the glassware in a hot air oven overnight, covering the mouths of the containers with aluminum foil.
- Make small holes in the aluminum foil using a syringe needle to allow for ventilation.
- 2.13. Finalize Cleaning:

- After drying, the glassware will be clean, dry, and sterile, ready for use in the synthesis of CDs.

### 7.3. Preparing Mango Leaf Extract

Materials Needed:20 ml ethanol, 10 ml Milli-Q water,2 grams mango leaf powder,50 ml bottle, Magnetic bead, stirring device,50 ml Falcon tube, Centrifuge

- 3.1. Mixing Solvent: In a clean 50 ml bottle, combine 20 ml of ethanol with 10 ml of Milli-Q water.
- 3.2. Incorporate Mango Leaf Powder: Introduce 2 g of mango leaf powder into the solvent mixture contained in the bottle.
- 3.3. Stir the Mixture: Place a magnetic bead into the bottle and stir the mixture for 4 hours at a speed of 400 rpm.
- 3.4. Clean the Bottle: Before starting, ensure that the bottle is cleaned with acetone.
- 3.5. Collect the Supernatant: After stirring, collect the supernatant from the mixture and transfer it into a 50 ml Falcon tube.
- 3.6. Centrifuge the Sample: Centrifuge the Falcon tube for 12-15 minutes at 8000 rpm.
- 3.7. Final Collection: After centrifugation, carefully collect the supernatant into the same or another 50 ml Falcon tube and discard the pellet. Your mango leaf extract is now ready for use.

### 7.4. Procedure for Reflux Set-up and Synthesis of mango leaves-derived quantum Dots (mQDs)

- 4.1. Prepare the Equipment:

- Take a round-bottom flask (RBF).
- Add the mango leaf extract and a magnetic bead into the RBF.
- 4.2. Set Up the Reflux System:

- Secure the RBF in place using Teflon tape on the condenser.
- Connect the outlet from the inlet pipe to the condenser.
- Ensure the entire RBF assembly is submerged in an oil bath without allowing it to touch the bottom of the flask.
- 4.3. Temperature Monitoring:

- Place a temperature probe in the oil bath.
- 4.4. Initiate the Reaction:

- Set all parameters on the hot plate magnetic stirrer.
- Start the reflux method, maintaining the following conditions:
- Temperature: 160 °C
- Speed: 400 rpm
- Reflux for a duration of 2 hours.
- 4.5. Cooling and Filtration:

- After a duration of 2 hours, let the solution reach room temperature.
- Allow it to cool, use a 0.22 µm filter to strain the solution and eliminate any particulates.
- 4.6. Evaporation:

- Use a rotary evaporator to evaporate the solvent.
- Set the rotary evaporator to 49 °C and 110 rpm.
- 4.7. Final Product:

- Collect the final product, which will be the carbon dots (CDs).
- 4.8. Store or Analyze:

- Store the synthesized CDs for further analysis or application as required. (Ensure to follow all safety protocols throughout the procedure.)

### 7.5. Characterization

**5.1 UV-Visible Spectroscopy:**

-1mg/ml mQDs stock solution, take 200 μL and dissolve in 2 mL Milli-Q water, and absorption is taken using UV-Vis spectroscopy.

-The result was further plotted using Origin software.

**5.2 Fluorescent spectrophotometer:**

-Prepare a 1 mg/ml stock solution of mQDs by taking 200 μL and dissolving it in 2 mL of Milli-Q water.

-Then, measure the fluorescence with a fluorescent spectrophotometer, and the results were plotted using Origin software.

**5.3. DLS and Zeta Potential:**

-The 1 mg/ml stock solution is filtered with a 0.22 μm filter. Take 200 μl of the filtered solution and dissolve it in 1 ml of Milli-Q water.

- Then they are subjected to the Malvern analytical Zetasizer Nano ZS instrument to measure the hydrodynamic size.

**Critical step**

To obtain precise size measurements, ensure there are no air bubbles in the cuvette during sample injection by utilizing a 1-ml syringe. If any air bubbles appear, gently tap the side of the cuvette to eliminate them. Additionally, any dust should be filtered out by using a 0.22-µm filter just before the measurement.

- The same instrument is used to measure zeta potential as well.

-The data is further analyzed using Origin software.

**5.4. Atomic Force Microscopy:**

-Cut a small piece of mica sheet and place it on a clean slide using clear nail paint adhesive. Let it dry.

-Prepare a 1:100 dilution of the 1 mg/mL mQDs stock solution in an MCT.

- Cleave the top layer of the mica sheet and drop cast the sample in a 1:10 dilution. Allow it to dry in a decorator.

- Image using AFM in tapping mode. Process the image using image software.

**5.5 Fourier Transform Infrared Spectroscopy (FTIR):**

-Scrap 1mg of the mQDs powder, perform FTIR analysis, and plot the data using Origin software.

**5.6 SEM (Scanning electron microscope)**

- Prepare a dilution from a 1 mg/ml solution by taking 200 μl and dissolving it in 2 ml of Milli-Q water. Drop the cast solution onto a 10 mm cover slide with a sample volume of 3-5 μl. Allow it to dry in a decorator.

- Image using SEM in Scanning mode.

-Process the image using image software.

**7.6. *In vitro:***

### 6.1 MTT Assay

-In the MTT experiment, RPE1 cells were grown in DMEM, while SUM-159A breast cancer cells were kept in HAMS-F12 media supplemented with 10% fetal bovine serum and antibiotics.

-cell types were maintained at 37 °C in a humidified incubator with 5% CO2. Following that, they were incubated in serum-free media for 15 minutes at the same temperature and CO2 conditions.

-The cells are seeded in a 96-well plate and allowed to grow until they reach 80% confluency. In each well, seed 1 × 10^4^ cells in 100 μl.

- Incubate for 24 h with complete media.

- Remove the media and prepare the mQDs treatment at concentrations of 100, 200, 300, 400, and

500 μg/ml in serum-free media for 24 hours.

- Take out serum-free media and prepare 0.5 mg/ml of MTT in serum-free media.

- add 100 μl to each well, incubate for 3-4 h

- Take out MTT media solution.

-Following the treatment, the medium was discarded, and 100 µl of MTT solution (0.5 mg/mL) was introduced. The cells were subsequently incubated for a duration of 3 to 4 hours.

-The absorbance was recorded at 565 nm utilizing a Byonoy’s Absorbance 96 microplate reader.

-Every experiment was conducted in sets of three, with untreated cells used as a control to assess the cell viability percentage for each well.

-The untreated mQDs acted as the control to evaluate the cell viability percentage for each well. The percentage of cell viability was determined using the following formula.

Formula:

Cell viability (%) = (Absorbance of the sample/Absorbance of the control) × 100.

### 6.2 Cellular Uptake

In the cellular uptake experiment, RPE1 cells were grown in DMEM, while SUM-159A (breast cancer) cells were kept in HAMS-F12 media.

**-** The cells were seeded in 4-well plates on a cover slip and allowed to grow until they reached 80% confluency. In each well. Seed each well 1 × 10^5^ cells. The well volume is 500 μl, and incubate for 24 h.

-Eliminate all media and rinse the seeded cells with 1× PBS buffer three times.

{Be aware of the PBS volume used for rinsing (450 μL)}.

-Subsequently, incubated in media without serum for 15 minutes at 37 °C with 5% CO2 in a humidified incubator.

-Discard the serum-free media and rinse three times with 1x PBS buffer.

-Following the washing step, the cells were exposed to modified quantum dots (mQDs) to evaluate their cellular internalization efficiency.

-Different concentrations (Control (no treatment), 50,100,200 μg/ml) were used for cell treatment. Allow the mixture to be incubated for 15 minutes at 37 °C with 5% CO2 in an incubator that maintains humidity.

- Remove treatment.

-The cells that were treated were fixed for 15 minutes at 37 °C using 400μl of 4% paraformaldehyde and were washed three times with 1× PBS.

-The cells were permeabilized using 0.1% Triton X-100 (300 μL) and subsequently stained with 0.1% phalloidin to visualize the actin filaments.

-Transferring Step (it is a little bit tricky) to take our coverslip from the well and fix it to the microscope slide.

-The cells were subjected to three washes with 1× PBS before being mounted onto the slides using Mowiol supplemented with 2.5 μl of DAPI for nuclear staining.

-Slide preparation: Wipe the slide with IPA thoroughly, and wipe with dry tissue per slide 2-4, cover the slide. Label the slides: cell line name, dates, and mount.

-The slides are stored at 4°C until imaging.

-Cell imaging was conducted utilizing a 63X oil immersion objective lens on a Leica SP8 laser scanning confocal microscope.

-The 405 nm laser is used for DAPI, and the 633 nm laser is used to analyze CDs.

-Quantitative image analysis was conducted utilizing Fiji, an advanced distribution of ImageJ software.

-The analysis entailed quantifying whole-cell intensity using maximum intensity projections, followed by background subtraction. Furthermore, the measured fluorescence intensity was normalized against that of unlabeled control cells to ensure accuracy in the assessment.

-A total of approximately 40-50 cells were analyzed from the acquired z-stacks for each experimental condition.

## 8. Result and Discussion

### 8.1. Characterization of QDs

To characterize the optical properties of Mango leaves-derived quantum dots (mQDs), we conducted an analysis using UV and fluorescence spectroscopy. The UV spectra revealed distinct peaks at 206 nm, attributable to the –OH group, and 260 nm, associated with flavonoids, as demonstrated in Figure (a). Upon excitation at various wavelengths (300, 350, 400, 460, 500, 560, and 600 nm), the mQDs exhibited maximum fluorescence intensity at 400 nm (Figure (b)). Additionally, Figure (c) illustrates the visual differences: mQDs exposed to daylight appear light yellow (left) while those under UV light emit a bright red fluorescence (right). Dynamic Light Scattering (DLS) was employed to ascertain the size of the mQDs, which, as depicted in Figure (d), show a predominant size of approximately 8.3 nm. The morphology of the mQDs was further examined via Atomic Force Microscopy (AFM), with results in Figure (e) indicating a quasi-spherical shape and a topographical size range of about 5 nm to 15 nm, as elaborated in Figure (h). Scanning Electron Microscopy (SEM) complemented our morphological analysis, revealing sizes between 10 to 18 nm and a consistently spherical shape, as shown in Figures (f) and (i). To probe the surface functionalization of the mQDs, we utilized Fourier Transform Infrared Spectroscopy (FTIR). The transmittance spectrum presented in Figure (g) compares the mQDs (black line) to mango leaf powder (mleaves) (red line) across a wavenumber range of 4000 to 500 cm⁻¹, with percentage transmittance indicated on the y-axis.

The mQDs showcased pronounced absorption bands corresponding to several functional groups: alcohol (–OH), sp³-CH stretch, carbonyl (C=O), alkene (C=C), ester (C=O), and C–O bonds, revealing a complex array of chemical functionalities. In contrast, the FTIR spectrum for the mango leaf powder exhibited a relatively featureless profile, indicating a limited presence of functional groups compared to the mQDs. This observation suggests substantial chemical modifications during the synthesis process of the mQDs from the leaf material.

### 8.2. MTT assay

Investigating the cytotoxicity of mQDs on RPE1 cells and SUM159A cells is essential for elucidating their biological impact. To assess this, an MTT assay was employed to quantify cell viability. The analysis was conducted across both cancerous (SUM159A) and non-cancerous (RPE1) cell lines to provide a comparative understanding of the mQDs’ effects. The different concentrations of mQDs (quantum dots) on the viability of RPE1 cells and SUM159A. The x-axis shows the concentration of mQDs in µg/mL (Control, 100, 200, 300, 400, 500), and the y-axis represents cell viability as a percentage. Figure 2(a) The results indicate that cell viability remains around or above 100% across all tested concentrations, with a notable increase in viability at the highest concentration (500 µg/mL), suggesting that mQDs are not cytotoxic to RPE1 cells within this range and may even promote cell growth at higher concentrations. Error bars are present, reflecting the variability in the data for each concentration tested. Figure (b) presents the bar graph showing the effect of mQDs on the viability of SUM159A cells at various concentrations (100 to 500 µg/mL), with the y-axis representing cell viability in per cent and the x-axis showing both control and treatment groups. The control group maintains near 100% cell viability, while all mQDs-treated groups show a marked decrease, with viability dropping to about 40–50% across all concentrations, indicating that mQDs have a pronounced cytotoxic effect on SUM159A cells regardless of dose within the tested range. Error bars are present for each group, showing variability, but overall, the trend demonstrates significantly reduced cell viability compared to the control.

**Figure 1.**
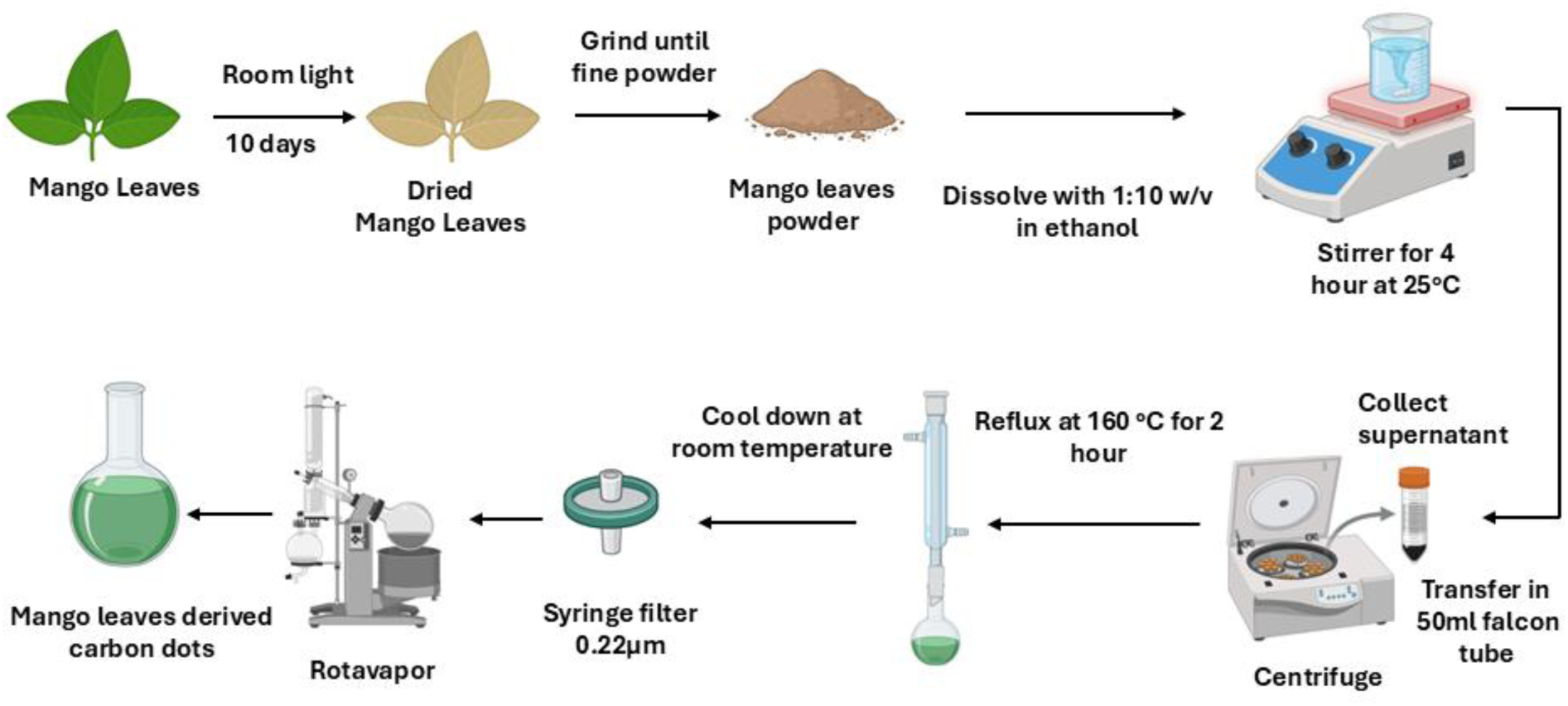
Synthesis of carbon dots involves using a top-down approach for synthesizing mango leaf-derived carbon dots. First, collect fresh mango. Clean and wash them thoroughly, then dry them at room temperature for 10-12 days. Once dry, grind the leaves into a fine powder. Dissolve the fine powder in ethanol at a 1:10 weight/volume ratio and immerse it for 4 hours at room temperature. Afterward, transfer the mixture to a Falcon tube and centrifuge it for 15 minutes at 8000 rpm. Collect the supernatant and discard the pellet. And prepare mango leaves extracts, take the mango leaf extract, and perform reflux carbonization at 160 °C for 2 hours while stirring at 400 rpm. Let the reaction mixture come down to room temperature, and afterward, filter it with a 0.22 µm filter. Evaporate the solution with a rotary evaporator to synthesize the mango leaf- derived carbon dots^21^. (Created in BioRender.Com).

**Figure 2.**
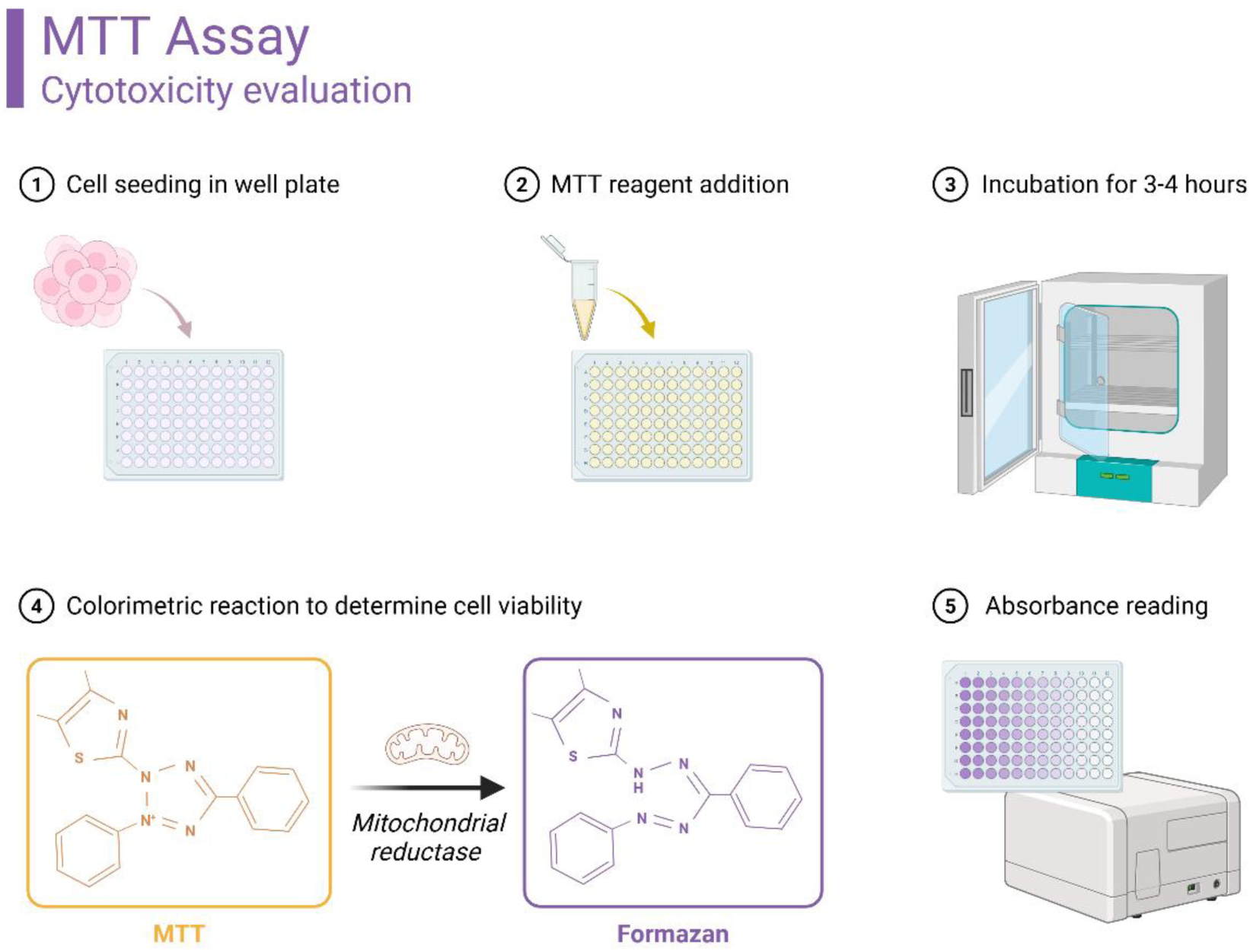
The MTT assay is a widely used method for assessing cytotoxicity and evaluating cell viability based on mitochondrial activity. 1st step: The process begins with cell seeding in a multi- well plate, where the cells are cultured. 2^nd^ step: Following this, a yellow MTT reagent is added to the wells.3^rd^ step: The plate is then incubated for 3 to 4 hours to facilitate the reaction. 4^th^ step: During this incubation, the yellow MTT compound is reduced by mitochondrial reductase present in viable cells, leading to the formation of purple Formazan crystals, which serve as an indicator of metabolically active cells. 5^th^ step: The final step involves reading the plate in a spectrophotometer or plate reader to measure the absorbance. The intensity of the purple colour correlates directly with the number of viable cells, allowing for the evaluation of cytotoxicity based on mitochondrial activity. (Created in BioRender).

### 8.3. Cellular Uptake

In this study, we explored the effects of modified quantum dots (mQDs) on various cell lines, with a particular emphasis on SUM-159A breast cancer cells due to their documented anti-cancer properties. We employed fluorescence signals emitted by mQDs alongside standard nuclear markers such as DAPI to evaluate cellular morphology and physiological responses. To determine the toxicity profiles of the mQDs, we conducted two sets of experiments involving both cancerous cells (document number 1) and non-cancerous epithelial cells (RPE-1). The cells were initially cultured and then exposed to increasing concentrations of mQDs. The results indicated a significant increase in uptake in both cancerous and epithelial cells at mQD concentrations of 100 and 200 μg mL−1, in contrast to the lower concentration of 50 μg mL−1 and the control group.

Consequently, cellular uptake assessments were standardized at concentrations of 100 μg mL^-1^ and 200 μg mL^-1^, as illustrated in Figures 3 and 4.

**Figure 3.**
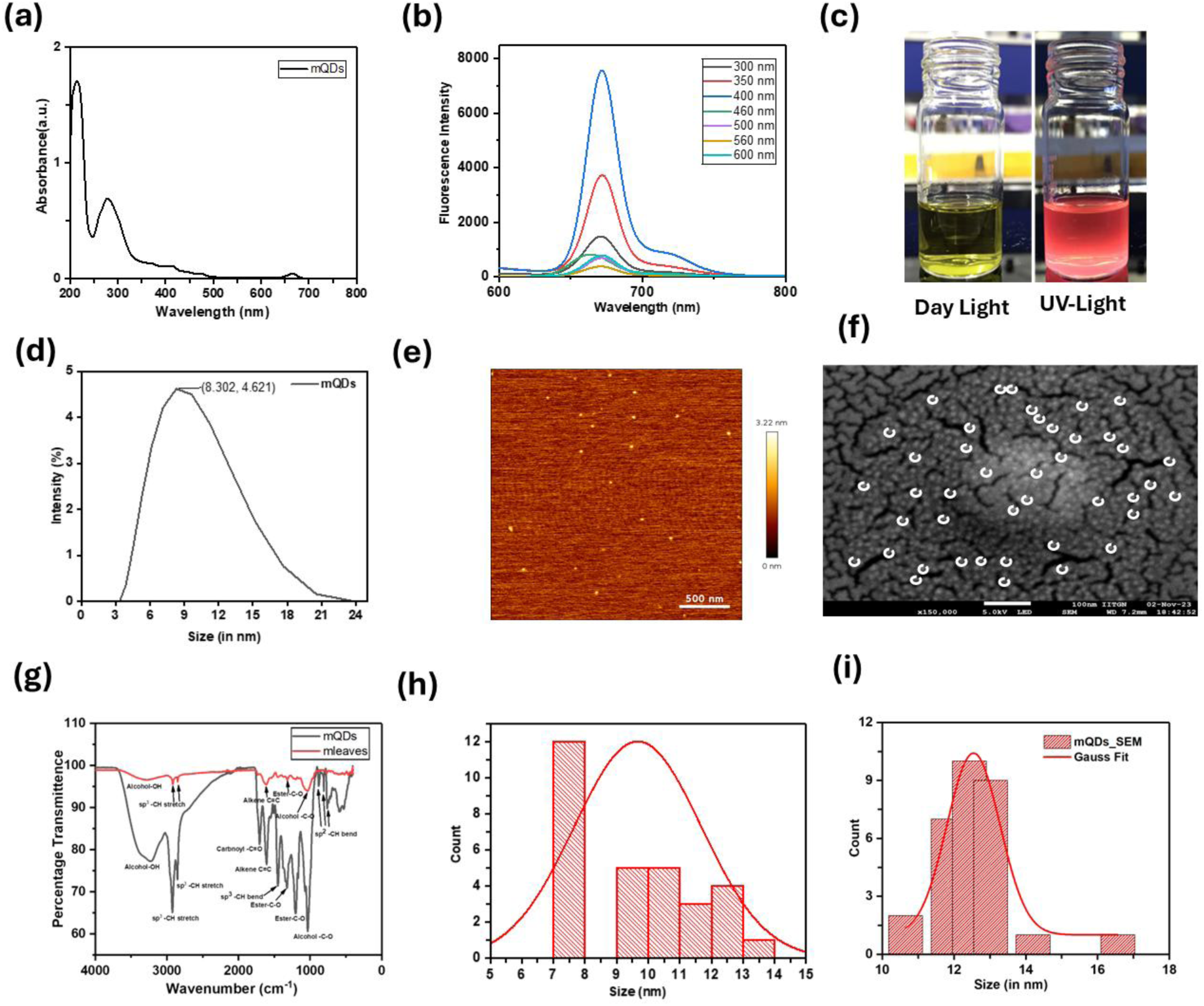
Characterization of mQDs: (a) The UV spectra of mQDs exhibited peaks at 206 nm (indicating the presence of –OH groups) and 260 nm (associated with flavonoids). (b) The mQDs were excited at various wavelengths (300, 350, 400, 460, 500, 560, and 600 nm), with the maximum intensity observed at a wavelength of 400 nm. (c) The fluorescence of mQDs is shown under daylight (left) and UV light (right). (d) Dynamic Light Scattering (DLS) analysis was conducted, (e) and (h), along with Atomic Force Microscopy (AFM) imaging and distribution of mQDs, (f) and(i), Scanning Electron Microscopy (SEM) to assess the morphology and size distribution. (g) Additionally, the FTIR spectrum was recorded.

**Figure 4.**
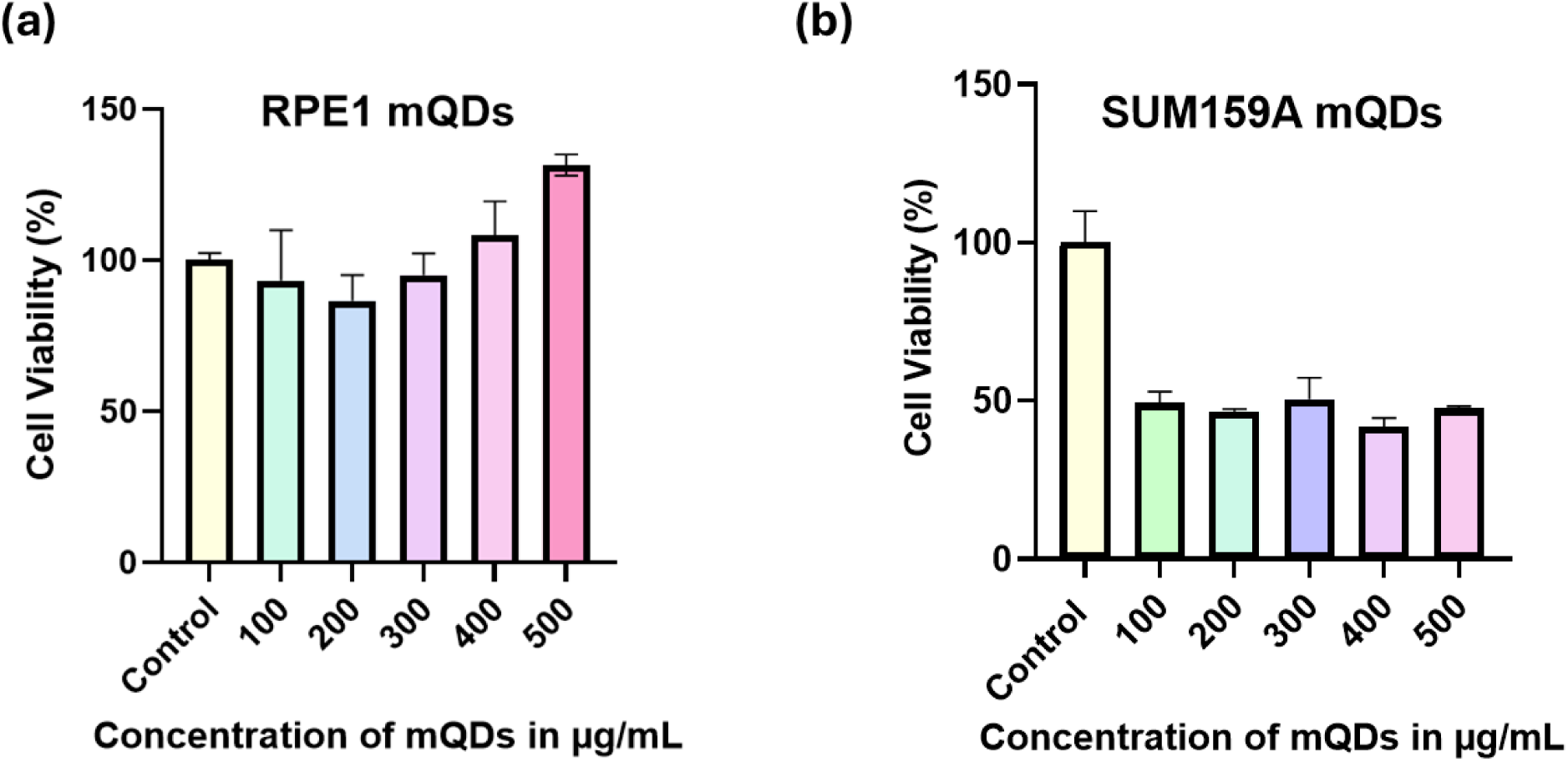
Cytotoxicity Assessment: (a) Evaluation of varying concentrations of mQDs (100, 200, 300, 400, and 500 μg mL−1) on RPE1 cells, and (b) assessment of the same concentrations of mQDs on SUM159A cells. The following statistical thresholds were applied to ascertain significance: ****p < 0.0001 indicates high significance, while ns (p > 0.05) denotes no significant effect.

**Figure 5.**
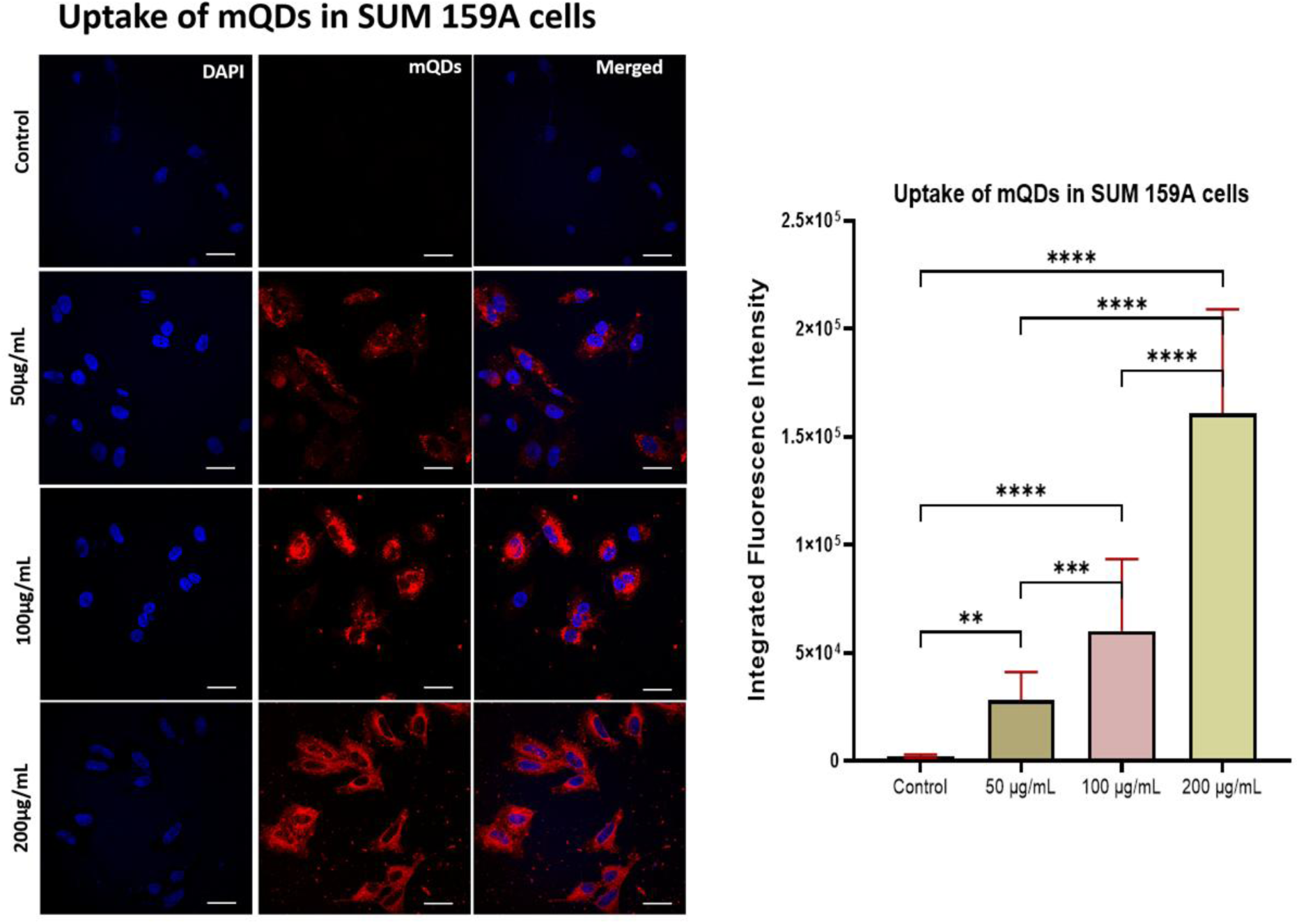
The cellular uptake of metal quantum dots (mQDs) in SUM-159A cells was examined, with a scale bar of 5 μm indicating measurement reference. Uptake was quantified at concentrations of 50 μg/mL, 100 μg/mL, and 200 μg/mL. Fluorescence intensity corresponding to the mQDs was measured within these cells. Statistical analysis was performed using one-way ANOVA in Prism Software, with significance indicated by **** for p < 0.0001 and ns for non- significant differences (n = 30).

**Figure 6.**
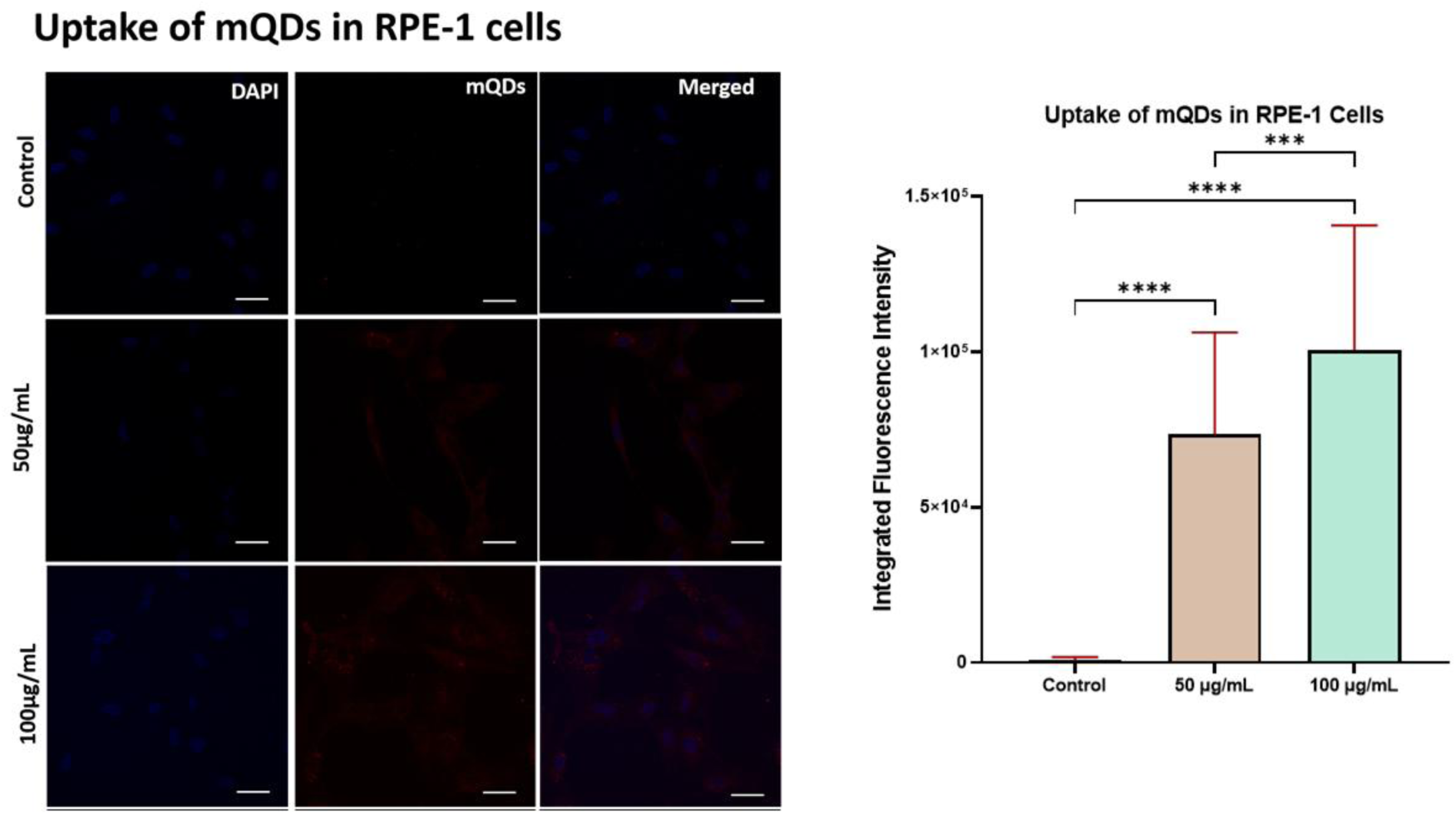
The uptake of mongo quantum dots (mQDs) in RPE-1 cells was evaluated, with a scale bar of 5 μm included for reference. Cellular uptake was assessed at concentrations of 50 μg/mL and 100 μg/mL. Fluorescence intensity of the mQDs within the RPE-1 cells was quantified, and statistical significance was determined using one-way ANOVA via Prism software. Results were reported as **** for p < 0.0001, with instances of no significant difference indicated as ns (n = 30).

## 9. Conclusion

In conclusion, the successful synthesis of red-emitting carbon quantum dots (mQDs) via a green synthesis approach presents a significant advancement toward environmentally friendly nanomaterials with promising applications in fluorescence-based technologies. Characterization of the mQDs revealed distinct spectral features, including peaks in UV spectra at 206 nm and 260 nm, indicating the presence of functional groups and flavonoids, respectively. The mQDs exhibited maximum fluorescence at an excitation wavelength of 400 nm and demonstrated favorable morphology and size distribution through various imaging techniques, including DLS, AFM, and SEM. The cytotoxicity assessment of mQDs on RPE1 non-cancerous cells showed remarkable cell viability, maintaining around or above 100% even at higher concentrations, suggesting a potential for mQDs to promote cell growth rather than induce toxicity. Conversely, the analysis of cancerous SUM159A cells demonstrated a pronounced cytotoxic effect, with cell viability dropping to approximately 40-50% across all tested concentrations. This disparity highlights the selective impact of mQDs, potentially opening avenues for targeted cancer therapies. Furthermore, the application of mQDs in fluorescence imaging, alongside markers like DAPI, has enabled insightful observation of cellular morphology. Together, these findings underscore the dual potential of mQDs as both harmless to healthy cells and effective against cancer cells, paving the way for future research into their therapeutic applications and contributions to sustainable nanotechnology.

## Data availability

Data supporting this study can be accessed in our earlier publications^2,22^ or requested from the corresponding author as needed.

## Supporting information

https://www.rsc.org/suppdata/d4/na/d4na00306c/d4na00306c1.pdf

https://pubs.acs.org/doi/suppl/10.1021/acsabm.4c00249/suppl_file/mt4c00249_si_001.pdf

